# CRISPR-induced exon skipping of β-catenin reveals tumorigenic mutants driving distinct subtypes of liver cancer

**DOI:** 10.1101/2022.03.27.485965

**Authors:** Haiwei Mou, Junjiayu Yue, Ying Jin, Zhikai Wang, Ya Gao, Tobias Janowitz, Hannah V. Meyer, Alper Kucukural, John E Wilkinson, Deniz M. Ozata, Semir Beyaz

**Affiliations:** Cold Spring Harbor Laboratory, Cold Spring Harbor, NY,11724; Bioinformatics Core, University of Massachusetts Medical School, 368 Plantation Street, Worcester, MA, 01605; Department of Comparative Medicine, University of Washington, Seattle, WA, USA; Department of Molecular Biosciences, The Wenner-Gren Institute, Stockholm University, S-106 91 Stockholm, Sweden

## Abstract

CRISPR/Cas9-driven cancer modeling studies are based on disruption of tumor suppressor genes (TSGs) by small insertions or deletions (indels) that lead to frame-shift mutations. In addition, CRISPR/Cas9 is widely used to define the significance of cancer oncogenes and genetic dependencies in loss-of function studies. However, how CRISPR/Cas9 influences gain-of-function oncogenic mutations is elusive. Here, we demonstrate that single guide RNA targeting exon 3 of *β-catenin* results in exon skipping and generates gain-of-function isoforms *in vivo*. CRISPR/Cas9-mediated exon skipping of *β-catenin* induces liver tumor formation in synergy with YAP^S127A^ in mice. We define two distinct exon skipping-induced tumor subtypes with different histological and transcriptional features. Notably, ectopic expression of two exon-skipped β-catenin transcript isoforms together with YAP^S127A^ phenocopies the two distinct subtypes of liver cancer. Moreover, we identify similar *β-catenin* exon skipping events in hepatocellular carcinoma (HCC) patients. Collectively, our findings advance our understanding of β-catenin-related tumorigenesis and reveal that CRISPR/Cas9 can be repurposed, in vivo, to study gain-of-function mutations of oncogenes in cancer.

## Introduction

CRISPR/Cas9 system is a powerful genetic tool that allows targeted genome editing (1, 2). CRIPSR/Cas9 introduces double strand breaks (DSBs) at targeted genomic locus through a programmed single guide RNA (sgRNA) and the Cas9 endonuclease (1, 2). Following CRISPR/Cas9-induced DSBs, small deletions or insertions (indels) are introduced by non-homologous end joining pathway which has been widely used for disruption of targeted genes (1).

Liver cancer is one of the most common cancers in the world and significantly contributes to cancer-related mortality (3, 4). The most frequent type of liver cancer in adults is Hepatocellular Carcinoma (HCC), which comprises more than 90% of all liver cancers (4). On the other hand, Hepatoblastoma (HB) is a rare form of liver cancer, which is predominant in children (5, 6). Aberrant oncogenic activation of Wnt/β-catenin signaling pathway is linked to both HCC and HB (4). Around 26% of human HCC patients present β-catenin mutations (7). Most of the hotspot mutations are within the exon 3 of *β-catenin* (8, 9), which encodes the domain containing phosphorylated serine/threonine residues targeting β-catenin for degradation in cytoplasm (10, 11). Once the phosphorylation site is mutated or deleted, degradation of β-catenin is impaired, thereby allowing unphosphorylated active β-catenin to accumulate, translocate into nucleus and to drive the transcription of various oncogenic target genes (10).

β-catenin mutations correlate with tumor progression (12) and have been used to refine liver cancer classification (13). In HB, different mutations of β-catenin associate with histologically and transcriptionally distinct subtypes (14). For instance, specific β-catenin mutations such as S45 show high β-catenin activity and is associated with hepatocellular adenoma (HCA) subtype (12). β-catenin mutations that influence its localization patterns also inform classification of HB subtypes (15). Thus, better understanding of β-catenin mutations in liver tumorigenesis is important to define disease heterogeneity and patient outcomes.

Mutated β-catenin alone is insufficient to form tumor in liver cells, but it drives tumorigenesis in synergy with other oncogenes like mutated Yap1 or c-Met (16, 17). Mutated β-catenin N90 (ΔN90) and YAP^S127A^ have been used to model liver cancer and are believed to drive tumors with features of HB (16). Depending on its oncogenic partner, β-catenin can also drive HCC (17). However, it is unknown whether different β-catenin mutations can drive different tumor types in synergy with the same oncogene. CRISPR/Cas9 gene editing is a common tool to model cancer *in vivo* (1, 2). Most of the CRISPR/Cas9-based cancer modeling involves the disruption of tumor suppressor genes (TSGs) by introducing indels that lead to frameshift mutations and loss of function (18-20). In this paradigm, CRISPR mediated indels of oncogenes lead to loss-of-functions. Whether indels also lead to gain-of-functions of oncogenes are not well known. We previously demonstrated that one sgRNA targeting exon 3 of *β-catenin* induces exon skipped in-frame transcripts in lung cancer Kras^G12D^;p53^−/–^ (KP) cell line (11). However, whether CRISPR-induced exon skipping occurs *in vivo* remains unknown.

Here, we deliver one sgRNA (sgCtnnb1) targeting exon 3 of *β-catenin*, together with Cas9 endonuclease, into liver cells by hydrodynamic tail vein injection (HDI). Immunohistochemical (IHC) staining reveals nuclear localization of active β-catenin upon injection. Moreover, simultaneous delivery of sgCtnnb1 and YAP^S127A^ confers growth advantage to exon skipped hepatocytes, thereby inducing tumor formation *in vivo*. Intriguingly, based on histology and transcriptome analyses, we report two distinct subtypes of liver tumor: one is induced by exon 3 deleted β-catenin isoform and YAP^S127A^, while the other is induced by YAP^S127A^ and β-catenin isoform whose exon 3 and 4 deleted. Furthermore, ectopic expression of two representative exon skipped β-catenin transcripts in synergy with YAP^S127A^ phenocopied our observed subtypes. Concordantly, two human HCC cases from The Cancer Genome Atlas reveal exon 3 and 4 deleted β-catenin isoform.

## Results

### sgRNA targeting *β-catenin* exon 3 induces nuclear accumulation of *β-catenin* in mouse liver cells

CRISPR-mediated gene editing approaches are commonly utilized to disrupt genes due to frame-shift mutations introduced by indels. However, how CRISPR affects Mrna splicing events and changes transcript isoforms are unknown. Here, we focused on the effect of CRISPR targeting exon 3 of *β-catenin* (*Ctnnb1*) on expression of different β-catenin isoforms. Exon 3 encodes phospho-acceptor residues that promotes the degradation of β-catenin (10). Therefore, genetic removal of *β-catenin* exon 3—that is in-frame with exon 4—blocks β-catenin degradation and leads to its active form that is trans-localized into nucleus (10, 11). While sgRNA targeting exon 3 of *β-catenin* (*Ctnnb1*) induces exon 3 skipped transcripts that are in-frame and produce gain-of-function nuclear β-catenin isoform in lung cancer KP cells *in vitro*, whether these CRISPR-induced isoforms occur *in vivo* is elusive (11).

To test whether CRISPR-mediated exon skipping events produce gain-of-function β-catenin isoforms *in vivo*, we utilized an sgRNA that targets *β-catenin* exon 3 (sgCtnnb1.1) (11) and cloned into px330 vector expressing SpCas9. We then delivered Cas9/sgCtnnb1.1 to liver cells via HDI (2) (Fig. 1A). We collected liver tissues from Cas9/sgCtnnb1.1 injected and non-injected mice at two weeks and two months post-HDI and performed RT-PCR to examine CRISPR-mediated exon skipping events. While we detected an additional ∼0.9 kb PCR band in injected liver compared to non-injected control liver at two weeks post-HDI, the band was not detected at two months post-HDI, suggesting that CRISPR-mediated exon skipping event in *β-catenin* takes place in low number of hepatocytes *in vivo* that are further replaced by non-edited hepatocytes (Fig. 1B). Consistently, IHC staining of β-catenin revealed that at two weeks post-HDI, 1.07% of hepatocytes (median = 1.07 ± 0.16%) retained nuclear β-catenin (Fig. 1C,1D) but at two months post-HDI, the positive staining of nuclear β-catenin was not detectable (Fig. 1C, 1D). Together, these data demonstrate that a single guide RNA targeting exon 3 of *β-catenin* is sufficient to induce a rare but an active exon 3-skipped β-catenin isoform which trans-localized into nucleus *in vivo*.

**Figure 1.**
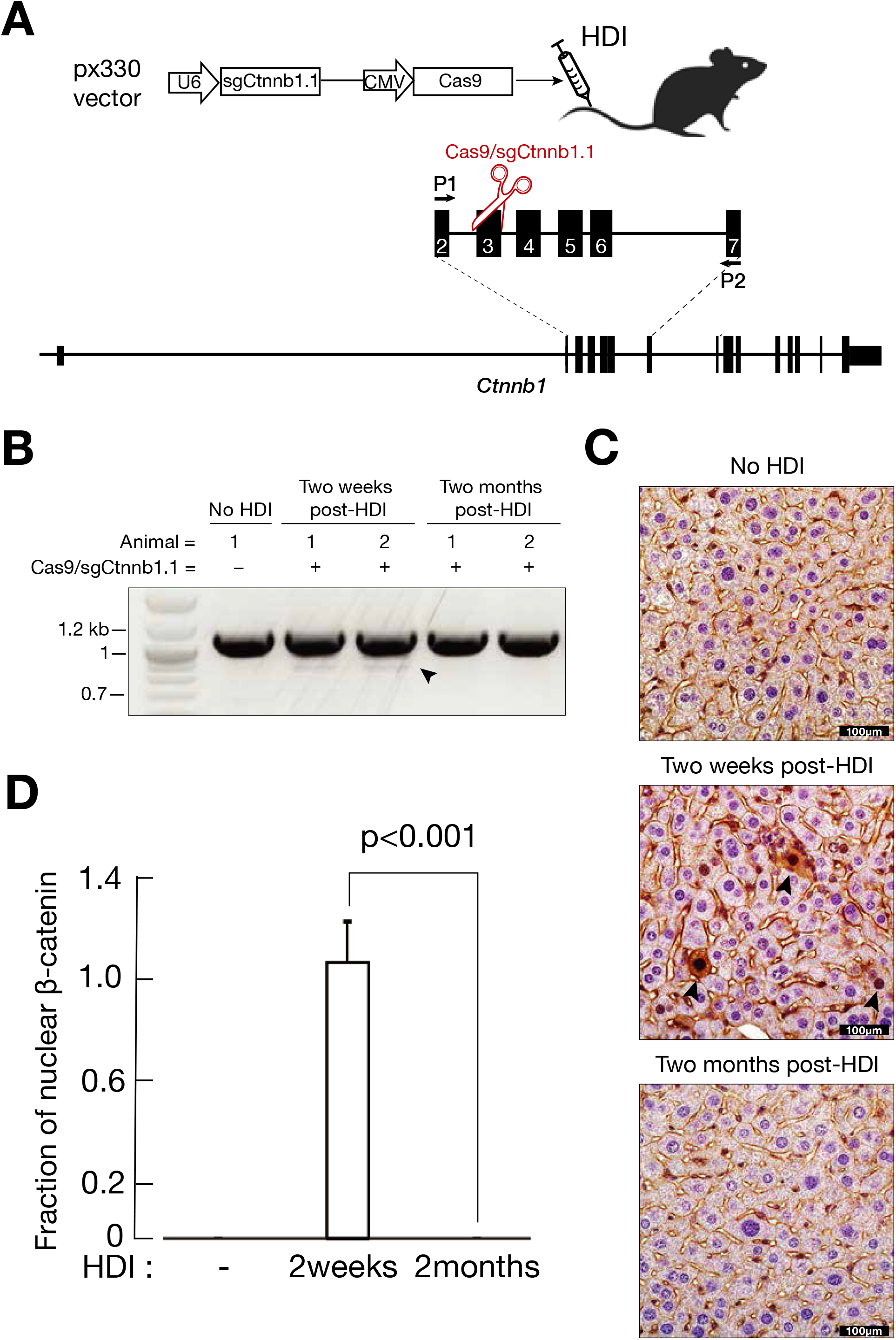
sgRNA targeting Exon 3 (sgCtnnb1.1) of *Ctnnb1* (β-catenin) induces skipped transcripts in mouse liver. (**A**) Schematic representation of hydrodynamic tail vein injection (HDI) that delivers sgCtnnb1.1 targeting Exon 3 of *Ctnnb1* to liver. P1 and P2 Primers were used to detect exon-skipped transcripts. (**B**) RT-PCR analysis to detect exon-skipped transcripts using primers, P1 and P2. (**C**) Immunohistochemical (IHC) detection of β-catenin from the liver tissues at two weeks and two months post-HDI. Non-injected liver tissue was used as negative control. Brown color shows β-catenin positive signal. (**D**) Cells with nuclear β-catenin and total hepatocytes were counted from six random fields of slides (two slides per mouse; two mice were counted). Positive cells divided by total cells to represent fraction of positive β-catenin nuclear staining. P value (p<0.001) indicates statistical significance.

### CRISPR-mediated *β-catenin* exon skipping along with YAP^S127A^ induces tumorigenesis in mice liver

CTNNB1 mutations are frequently observed in HB patients, especially in exon 3 and exon4 (Fig. 2A) (6). Moreover, activation of β-catenin together with YAP^S127A^—mutated YAP that is constitutively active and remains in the nucleus—induces hepatoblastoma (16). In line with this, we hypothesized that if CRISPR-mediated exon skipping generates a functional nuclear β-catenin, and cells with both exon-skipped β-catenin and YAP^S127A^ will have a growth advantage and form liver tumor.

**Figure 2.**
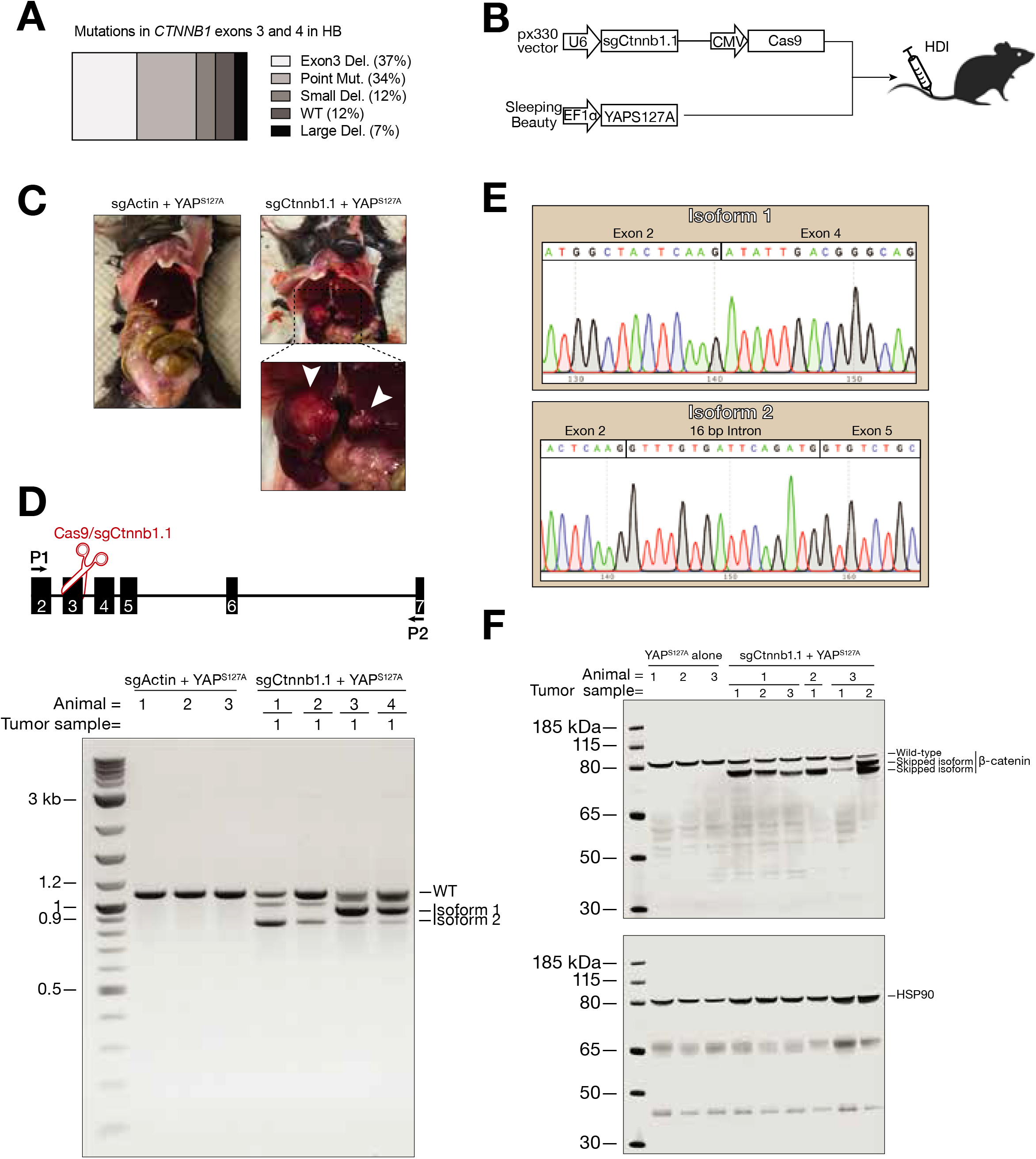
YAP^S127A^ confers exon skipping hepatocytes with tumorigenic ability. (**A**) Mutation frequency of CTNNB1 Exon 3 and 4 in HB. (**B**) Schematic representation of hydrodynamic injection (HDI) that delivers Sleeping-Beauty (SB) based YAP^S127A^ and sgRNA targeting Exon 3 of Ctnnb1, sgCtnnb1.1. (**C**) Representative necropsy photographs of tumor modeled by YAP^S127A^ and sgCtnnb1. Arrows indicate tumors. (**D**) Total RNA isolated from tumors were subjected to RT-PCR to detect wild-type (WT) or exon skipping bands. P1 and P2 Primers were used as forward and reverse primers. (**E**) Sanger sequencing showing two exon-skipped transcripts: isoform 1 where entire Exon 3 skipped and isoform 2 where Exon 3 and 4 fully skipped but with 16 nucleotide intron insertion. (**F**) Western blotting shows the wild-type and β-catenin skipped proteins from tumor samples (*n* = 10) of YAP^S127A^ and sgCtnnb1.1 injected mice (*n* = 4) and from liver tissues of YAP^S127A^ injected control mice (*n* = 3). HSP90 is used as a loading control.

To directly test our hypothesis, we delivered the Sleeping Beauty-based YAP^S127A^ (SB-YAP^S127A^) and CRISPR-based sgCtnnb1.1/Cas9 together to liver cells via HDI (Fig. 2B). At two months post-HDI, we observed gross tumor formation in liver (Fig. 2C). As a control, YAP^S127A^ and CRISPR-based sgActin/Cas9 showed healthy and tumor-free liver (Fig. 2C). To further examine whether the tumors collected at two months post-HDI are enriched for exon skipping *Ctnnb1* isoforms, we isolated total RNA from freshly frozen tumors and performed RT-PCR to detect exon skipped isoforms. In liver tumor samples, we detected two major *Ctnnb1* skipping isoforms (Fig. 2D). We then Sanger sequenced these two bands: isoform 1 skipped entire exon 3 that is in-frame with exon 4; isoform 2 skipped entire exon 3 and exon 4 but with 16 nucleotides intronic sequence, making this isoform be in-frame with Exon 5 (Fig. 2E). To test whether these skipping *β-catenin* isoforms encode proteins, we performed western blotting and observed that additional β-catenin protein bands appeared in tumor samples with smaller molecular weight compared to control liver tissues (Fig. 2F).

To rule out CRISPR off-target effect, we designed two more sgRNAs targeting *β-catenin* exon 3 with different protospacer adjacent motif (PAM) sequences. In concordance with sgCtnnb1.1, these two different sgRNAs (sgCtnnb1.2 and sgCtnnb1.3) induced tumor formation in synergy with YAP^S127A^ (Fig. S1A). Our IHC staining of β-catenin and YAP revealed positive staining of both proteins in liver tumor cells and confirmed that CRISPR-mediated exon 3 skipping of *β-catenin* produces gain-of-function β-catenin that in turn promotes tumor formation along with YAP^S127A^ (Fig. S1A).

To further substantiate that CRISPR-mediated targeting of *Ctnnb1* exon 3 produces gain-of-function β-catenin that promotes tumor formation *in vivo*, we delivered another oncogene, *c-Met*, alongside sgCtnnb1.1 to liver cells via HDI (Fig. S1B). At one-month post-HDI, we collected liver tissue and performed histological analysis and found tumor formation (Fig. S1B). Together, these results demonstrate that a single guide RNA targeting exon 3 of β-catenin results in exon skipped isoforms that act synergistically with oncogenes such as YAP^S127A^ or *c-Met* to drive liver tumor formation.

### Exon skipped β-catenin isoforms and YAP^S127A^ induce histologically and transcriptionally distinct liver tumors in mice

To further characterize liver tumors induced by the combination of CRISPR-mediated exon skipped β-catenin and YAP^S127A^, we examined the tumor sections and encountered two histologically distinct tumor lesions which we henceforward call subtype A and subtype B (Fig 3A). Subtype A is characterized by pleomorphic neoplastic hepatoblasts with particularly variable sized round to oval nuclei (anisokaryosis) and uniformly condensed chromatin. The neoplastic cells nuclear/cytoplasmic ratio varies significantly. There is extensive cytoplasmic clearing, especially perinuclear, and occasional mild accumulation of micro-vesicles. These cytoplasmic changes are reminiscent of glycogen and lipid accumulation associated with altered metabolism. Subtype B tumors are characterized by uniform small neoplastic epithelial hepatoblasts with relatively small, round nuclei with conspicuous peripheral clumping of chromatin and central pallor. There is mild cytologic atypia and occasional nuclei demonstrate prominent nucleoli. The cells have eosinophilic cytoplasm. For both subtypes, occasional mitotic figures are present, and the neoplastic masses are not encapsulated but are well demarcated with mild compression of the adjacent liver (Fig. 3A). Next, to assess lipid accumulation in these histologically distinct tumor subtypes, we performed Oil Red O staining that stains for neutral triglycerides and lipids on frozen sections. We found that the cytosol of subtype A cells showed clear Oil Red O staining, while the signal was below the limit of detection in the cytosol of subtype B cells (Fig. 3B). Interestingly, while both subtypes A and B showed similar YAP nuclei positive signal, the localization pattern of β-catenin was different between subtypes. Specifically, while subtype B showed the expected nuclear β-catenin staining pattern of active β-catenin isoforms, subtype A tumors surprisingly did not exhibit a clear nuclear β-catenin accumulation. Instead, β-catenin was detected mostly around the cytoplasm and membrane in subtype A tumors (Fig. 3A). Both subtype A and B tumors showed Ki67 positive staining from IHC staining, which is a well-established proliferation marker for tumors (Fig. S1C). These data indicate that CRISPR-mediated exon skipping results in oncogenic isoforms of β-catenin that drive histologically distinct liver tumors in mice and exhibit different cellular localization patterns.

**Figure 3.**
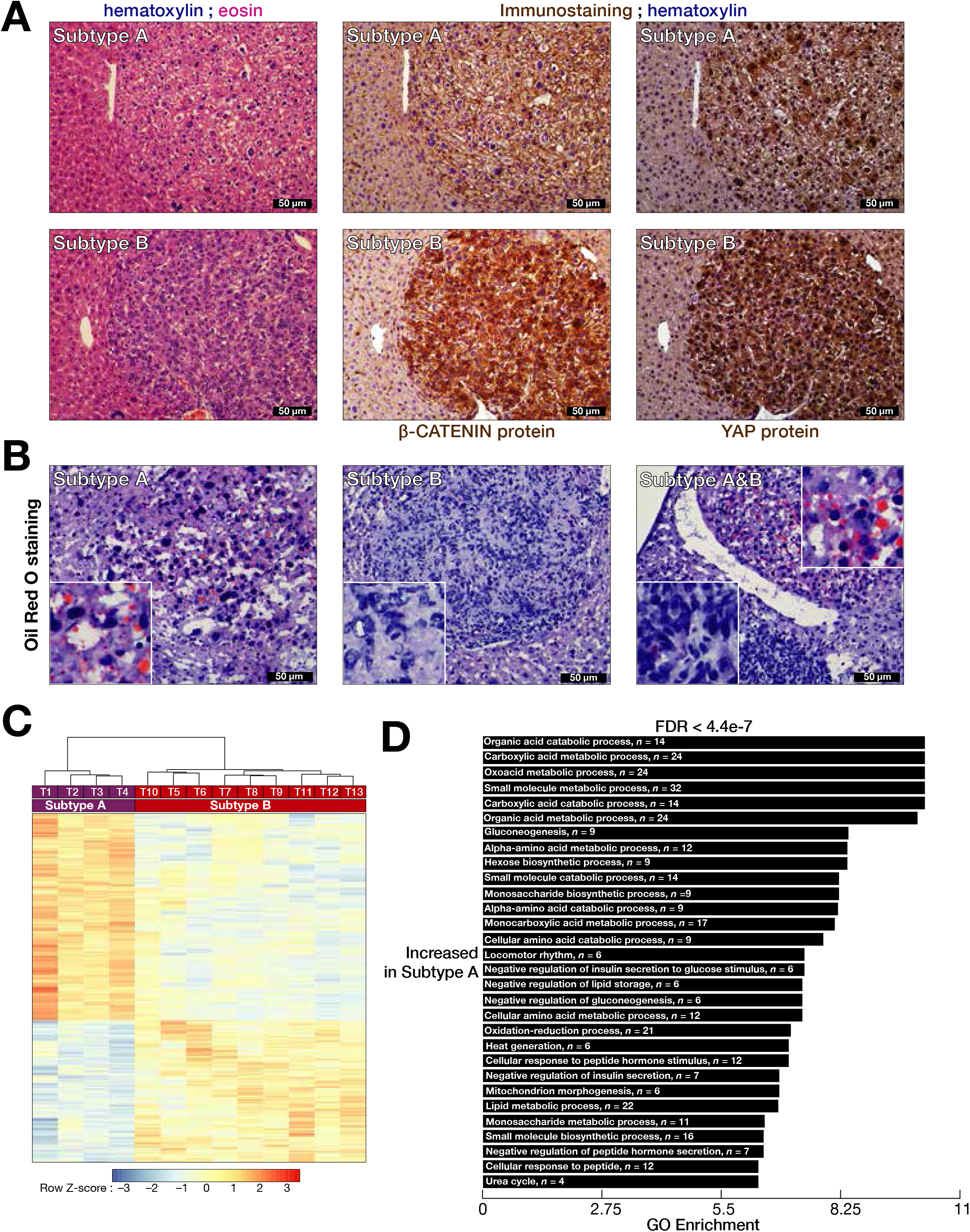
Histology and transcriptome analyses show two distinct subtypes of liver cancer. (**A**) Hematoxylin and eosin (H&E) staining shows the histology of Subtype A and Subtype B, while immunohistochemistry (IHC) shows β-catenin and YAP protein staining from these two subtypes modeled by sgCtnnb1 and YAP^S127A^. (**B**) Oil Red O staining shows lipid accumulation within the cytoplasm of two subtypes of liver tumor. (**C**) Supervised hierarchical clustering obtained from 3,347 differentially expressed genes shows distinct transcriptome signature of Subtype A and B. (**D**) Gene Sets Enrichment Analysis (GSEA) based on 1,979 up-regulated genes from Subtype A tumors shows metabolic pathway enrichment including lipid metabolism.

To determine the transcriptional signatures of these histologically distinct tumor subtypes driven by exon skipped β-catenin isoforms, we randomly collected 13 individual liver tumor samples and three normal liver tissue samples from five different mice and performed RNA sequencing. Unsupervised hierarchical clustering of the all 13 liver tumor samples revealed 4,800 differentially expressed genes in tumors compared to normal liver (<2-fold and >2-fold change, FDR < 0.01). Expression levels of well-established liver tumor marker genes, such as *Gpc3, Mki67, Afp* and *Cd34* (21-26), were increased in tumor groups compared to control (Fig. S1D). Clustering analysis identified two major groups of tumor samples: T1 to T4 were in one group (subtype A), while T5 to T13 were in another group (subtype B) (Fig. S2A). Differential gene expression analysis between the two groups of tumor samples revealed 3,347 differentially expressed genes (<2-fold and >2-fold change, FDR < 0.01) (Fig. 3C). To define functional categories enriched in distinct subtypes, we performed Gene Set Enrichment analysis (GSEA). Consistent with histological features of subtype A, metabolic pathways, including lipid metabolism were among the top functional categories enriched in upregulated genes in T1 to T4 tumor samples (subtype A) (Fig. 3D). Pathways that were enriched in upregulated genes for T5 to T13 (subtype B) consist of tissue development and morphogenesis (Fig. S2B). Importantly, our GSEA confirmed the liver tumor signature and YAP activation signature of all 13 tumor samples compared to normal liver (Fig. S2C). Because these subtypes exhibited distinct β-catenin localization patterns, we compared the expression levels of β-catenin target genes between subtype A and B. Several notable β-catenin targets were differentially expressed including *Id2, Lgr4, Cd44, S100a6* (Fig. S3). Together, our data indicate that exon-skipped β-catenin isoforms synergize with YAP^S127A^ to drive two histologically and transcriptionally distinct tumor subsets exhibiting differences in lipid metabolism and expression of β-catenin target genes. How different β-catenin isoforms influence lipid metabolism and β-catenin target gene transcription in these two distinct tumor subtypes warrants further investigation.

### Ectopic expression of exon skipped β-catenin isoforms together with YAP^S127A^ drive distinct liver tumor subtypes

To definitively assess whether the distinct subtypes of tumors we observed are driven by the oncogenic effect of CRISPR-induced exon skipped β-catenin isoforms rather than the off-target activity of Cas9 endonuclease inducing random mutations at other genomic sites, we cloned the two in-frame cDNA of skipped β-catenin isoforms into Sleeping Beauty system and delivered together with YAP^S127A^ into liver cells by HDI: one with entire exon 3 skipped (Skip-1), and the other with both exon 3 and exon 4 skipped entirely but with 16 nucleotides of intron insertion (Skip-2). Introducing both β-catenin isoforms along with YAP^S127A^ into liver cells formed tumor (Fig. 4A). To validate that Skip-1 and Skip-2 β-catenin isoforms are in-frame to produce β-catenin proteins, we performed western blotting from the tumor protein lysates and identified the expected molecular weight of Skip-1 and -2 isoforms (Fig. 4B).

**Figure 4.**
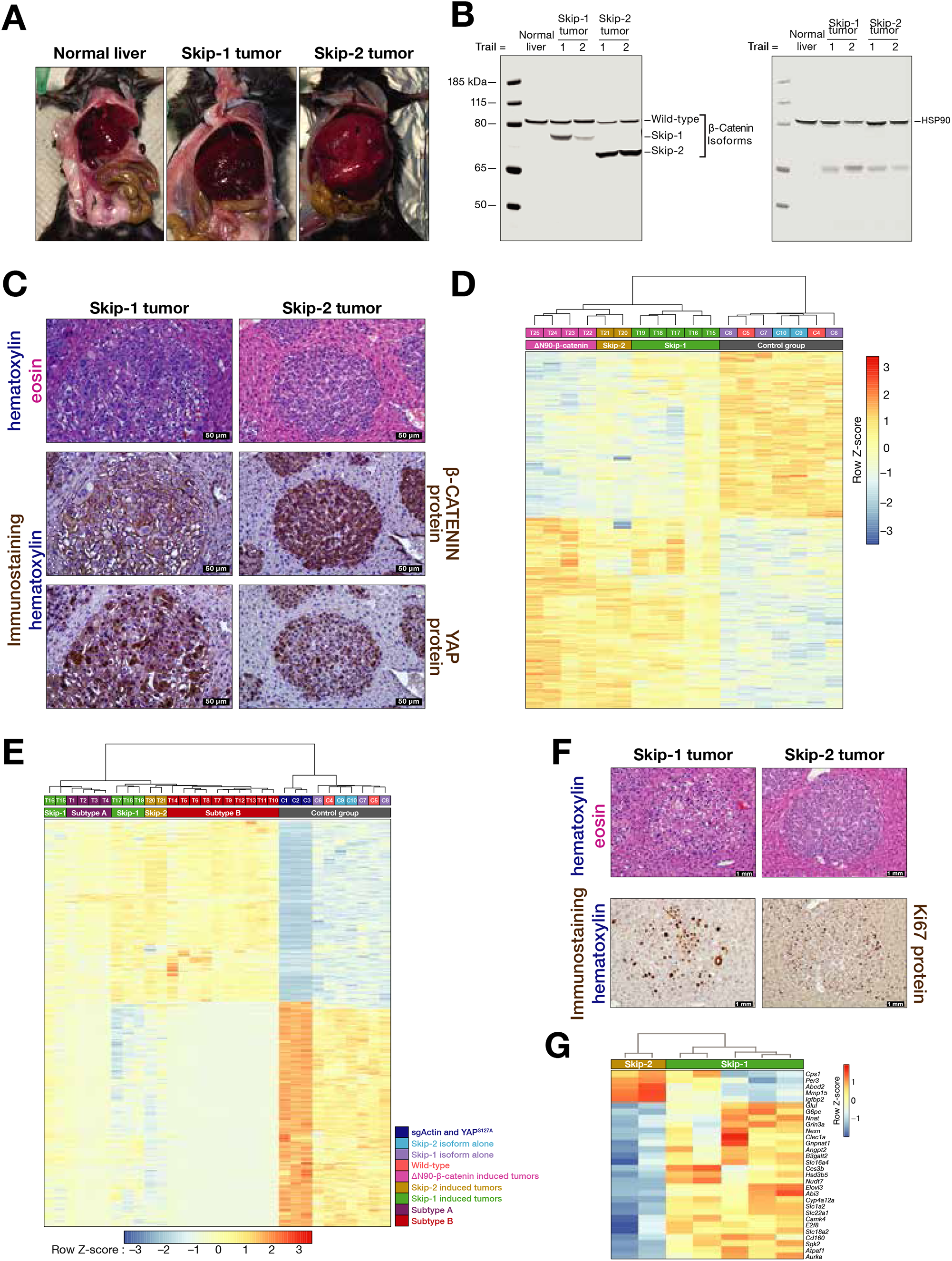
Ectopic expression of two exon skipped β-catenin transcripts show tumorigenicity in synergy with YAP^S127A^. (**A**) Representative tumor images induced by ectopic expression of Skip-1 or -2 transcripts in synergy with YAP^S127A^. (**B**) Western blotting shows the wild-type and β-catenin skipped proteins from tumor samples (*n* = 2) of Skip-1 and -2 induced tumors. Normal liver served as a negative control. HSP90 is used as a loading control. (**C**) H&E staining shows the histology of Skip-1 and -2 driven tumors. IHC shows β-catenin and YAP protein staining from Skip-1 and -2 driven tumors. Brown color indicates positive IHC signal. (**D** & **E**) Unsupervised hierarchical clustering groups Skip-1, Skip-2 and ΔN90-β-catenin driven tumors or Subtype A, Subtype B, Skip-1, Skip-2 and ΔN90-β-catenin driven tumors based on their transcriptome signature. (**F**) H&E and Ki67 IHC stainings from the serial sections of Skip-1 and Skip-2 driven tumors. Brown color indicates positive Ki67 IHC signal. (**G**) Heatmap shows highlighted differentially expressed genes related to lipid metabolism or β-catenin target genes between Skip-1 and Skip-2 driven tumors.

To further characterize tumor histology, we performed H&E staining. Gross histology of tumor samples revealed that similar to subtype A, Skip-1-induced tumor cells retained larger nuclei and unstained cytosol (Fig. 4C). On the contrary, similar to subtype B, Skip-2 induced tumor cells showed smaller nuclei and more structured and condensed tumor cells which is typically the histological hallmark of hepatoblastoma (16) (Fig. 4C). Immunohistochemistry staining showed that both tumors express YAP and β-catenin (Fig. 4C). In line with what we observed in subtype A, Skip-1 induced tumors retained less nuclear staining for β-catenin compared to Skip-2 (Fig 4C).

To determine the transcriptional profile of the tumors that are induced by Skip-1 and Skip-2 isoforms, we performed RNA sequencing. Using whole transcriptome, our principal component analysis (PCA) distinguished Skip-2 tumors from Skip-1 tumors and located Skip-2 tumor samples near ΔN90-β-catenin/YAP^S127A^ induced hepatoblastoma tumor samples, an established hepatoblastoma model (16)(Fig. S4A). Furthermore, control group mice that were injected with either Skip-1 or -2 isoforms without YAP^S127A^ were located near wild-type animals (Fig. S4A). In line with the PCA, unsupervised hierarchical clustering analysis grouped Skip-2 induced tumors with ΔN90-β-catenin induced tumors, while skip-1 induced tumors were separated from them (Fig. 4D).

Given the histology of Skip-1 induced tumors resemble subtype A and Skip-2 induced tumors resemble subtype B, we further examined whether the transcriptome of Skip-1 and -2 induced tumors is similar to that of subtype A and B tumors. We, therefore, performed unsupervised hierarchical clustering of all tumor samples along with control groups using 4,800 genes that were differentially expressed in subtype A and B compared to their corresponding controls. We found that Skip-2 induced tumors were clustered with subtype B, while Skip-1 induced tumors were clustered with subtype A (Fig. 4E). Moreover, Gene Ontology enrichment analysis using differentially expressed genes in Skip-1 and Skip-2 compared to their corresponding controls revealed that only Skip-2 induced tumors retained hepatoblastoma signature as evidenced by enrichment in hepatoblastoma pathway genes and that both Skip-1 and Skip-2 tumors showed YAP activation signature (Fig. S4B). Of note, both Skip-1 and Skip-2 tumors showed Ki67 IHC positive staining, indicating active proliferation of tumor cells (Fig. 4F). Moreover, the mRNA abundance of liver tumor marker genes, *Cd34, Gpc3* and *Mki67* (21, 22, 25, 26) were increased in both Skip-1 and Skip-2 tumors compared to controls (Fig. S4C). Finally, among differentially expressed genes between Skip-1 and Skip-2 tumors, several β-catenin pathway genes such as *Cps1, Abcd2, Mmp15*, and *Igfbp2* (27-30) were upregulated in Skip-2 tumors, whereas metabolic genes including the fatty acid elongase *Elovl3* were upregulated in Skip-1 tumors (Fig. 4G). These results demonstrate that ectopic expression of exon-skipped β-catenin isoforms give rise to distinct liver tumor subtypes with different histological and transcriptional features.

Because one sgRNA (sgCnnb1.1, sgCtnnb1.2 or sgCtnnb1.3) targeting exon 3 of *β-catenin* induces exon 3-skipped functional β-catenin isoform that acts along with YAP^S127A^ and leads to tumor formation, we tested whether using two sgRNAs that targets introns before and after exon 3 also produces gain-of-function β-catenin isoform *in vivo* and induces tumor formation in the presence of YAP^S127A^ (Fig. S5A). We first tested the genome editing capacity of our newly designed sgRNAs targeting introns above in KP cells: our RT-PCR analysis revealed ∼0.9 kb long PCR band (Fig. S5B) encoding a shorter protein of β-catenin skipped isoform (Fig. S5C), confirming that removing exon 3 is in frame with exon 4. We next delivered these two sgRNAs along with YAP^S127A^ to liver cells via HDI to test the tumorigenic capacity of the exon 3 deleted β-catenin isoform. At two months post-HDI, we observed gross tumor formation (Fig. S5D). Histological analysis of these tumors revealed consistent features with subtype A and Skip-1 induced tumors (Figs. 3A and 4C)—both were driven by the removal of exon 3 of β-catenin (Fig. S5E). These data indicate that loss of β-catenin exon 3 using an alternative targeting strategy results in oncogenic isoform that drives liver tumor formation *in vivo*.

### Exon 3 and 4 deleted β-catenin isoform is found in human hepatocellular carcinoma

We next utilized a HCC model that is driven by activation of β-catenin in combination with c-Met. To further test whether skipped β-catenin isoform recapitulates this HCC model, we delivered sleeping-beauty based cMet and CRISPR based β-catenin exon skipping, and as expected, we observed tumor formation (Fig. S6A and S6B), suggesting that exon-skipped β-catenin not only drive tumorigenesis with active YAP to recapitulate HB model, but also with c-Met for HCC model.

Given we identified two different CRISPR-mediated isoforms of β-catenin leading to histologically distinct liver tumor in mice, we next sought to understand whether these β-catenin isoforms are present in the etiology of human liver cancer. Therefore, we analyzed the RNA sequencing data of human tissue and cancer samples from The Cancer Genome Atlas (TCGA) for these exon skipping events. Our analysis captured exon skipping events in all analyzed cases. Although our analysis did not capture exon 3 deleted β-catenin isoform, intriguingly, two human hepatocellular carcinoma (HCC) cases revealed β-catenin isoform with the exclusion of exon 3 and 4 (similar to Subtype B and Skip-2 isoforms in mice), compared to other HCC cases (Fig. S6C and S6D).

## Discussion

Herein we report that a sgRNA targeting exon 3 of *Ctnnb1* (β-catenin) results in exon-skipped transcripts *in vivo*. These exon-skipped isoforms of β-catenin encode functional proteins that act synergistically with other oncogenes, YAP^S127A^ or c-Met to induce liver tumor formation. More importantly, CRISPR-mediated exon skipping events generate two distinct β-catenin isoforms that drive two discrete liver tumor subtypes.

### Exon skipping and alternative splicing

Although CRISPR-mediated genome editing driven by a sgRNA has been used to disrupt gene activity, we recently reported in lung adenocarcinoma cell line that one sgRNA can generate large deletions or alternative splicing in exons of oncogenes triggering aberrant juxtaposition of exons that encode shorter protein isoforms (11). The exact underlying mechanism leading to CRISPR-induced exon skipping remains elusive. Mutations affecting splice-sites can lead to alternative splicing events (31, 32). Accordingly, the indels within the exon 3 of *β-catenin* introduced by CRISPR leading to the disruption of exonic splicing enhancer sites could be a potential contributing factor for the complete skipping of exon 3. Alternatively, if splice sites in exon 3 and 4 are weak, the DNA insertion meditated by CRISPR can create stronger cryptic splice sites in the flanking intron that can lead to the inclusion of intronic fragment, while exon 3 and 4 are removed. A recent study suggests that the disruption of essential splice sites by premature termination codon mutations may be responsible for exon skipping (33).

Moreover, altered chromatin modifications (34), posttranslational regulation of the exon skipping machinery (35) or mRNA regulating proteins like polypyrimidine tract-binding proteins 1 (PTBP1) are other potential mechanisms underlying exon skipping (36)

The entire loss of exon 3 of *β-catenin* is recently reported in microsatellite stable (MSS) colorectal cancer (CRC) (37, 38). Interstitial deletions involving exon 3 induces oncogenic β-catenin in primary colorectal carcinomas without APC mutations (Iwao et al. Cancer Research 1998). In addition to the exon skipping events within the oncogenic *β-catenin* gene, these events are also observed in other oncogenes, like BRCA1 and MET (39, 40) in other cancers (41, 42). More importantly, exon skipping or splicing has already taken as a therapeutic tool (43, 44). There are studies showing the significance of exon skipping in other disease settings as well, such as altered splicing results in in-frame deletion of exons of *COL11A2, FBN1, FLOT1* genes (45-47). Together, utilizing CRISPR-based exon skipping may provide invaluable tool to model disease *in vivo* to better understand the underlying molecular mechanisms. We also envision that CRISPR-mediated exon skipping phenomenon may be used as a therapeutic tool to correct diseases. For example, Dushenne Muscular Dystrophy patients retaining exon 44 reveal more severe disease progression (48).

Our application of CRISPR-mediated exon skipping yields gain-of function β-catenin isoforms *in vivo*. Our data clearly indicate that introducing indels by a single sgRNA not only leads to loss of function, but also efficiently leads to gain-of function mutations. We, therefore, recommend that more careful CRISPR genotyping should be employed, as it may lead to the unexpected discovery of a potential disease-linked gain-of function mutation.

### CRISPR-induced β-catenin isoforms drive distinct liver tumor subtypes

We unexpectedly found two histologically distinct tumors driven by alternatively spliced β-catenin isoforms (Figs. 2 and 3). Tumor cells induced by Skip-1 isoform with whole exon 3 deletion showed lipid accumulation within their cytosol and reduced nuclear β-catenin, while Skip-2 induced tumor cells revealed absence of lipid accumulation in their cytosol and strong nuclear β-catenin staining.

Mutations in exon 3 of β-catenin correlate with β-catenin accumulation patterns in HCC. While consistent with this, our transcriptome analysis revealed differential expression of β-catenin target genes (Fig. 4G). Particularly, our analysis demonstrated that the mRNA abundance of *Glul*, classical β-catenin target gene, was increased in Skip-1 induced tumors, despite the lack of nuclear β-catenin compared to Skip-2 induced tumors. Concordantly, two subsets of human hepatocellular carcinoma patients both bearing activated β-catenin show different pattern of β-catenin IHC staining, one subset with nuclear β-catenin signal while the other without nuclear signal (49). Moreover, HCC patients with β-catenin exon 3 in-frame deletion exhibit increased *GLUL* expression, despite they retained less nuclear β-catenin staining (12), suggesting that β-catenin target genes are activated through additional events.

Our findings also provide insight into pathways downstream of β-catenin, potentially opening a window for identifying new physical interaction partners of β-catenin and the dynamics of how β-catenin interacts with other oncogenes to drive distinct tumor subsets. This model can be expanded to studies regarding the functional domains of proteins since skipping transcripts might generate potential gain-of-function mutations. Apart from cancer modeling, this exon skipping model can also be applied to and advance our understanding of alternative splicing *in vivo*.

## Methods

### Plasmids and cloning

Sleeping beauty based β-catenin N90 (ΔN90) (Addgene; 31785), YAP (Addgene; 86497), and c-Met (Addgene; 31784) plasmids were purchased from Addgene. gBlocks Gene Fragments of skipped β-catenin isoforms for Gibson cloning were synthesized by Integrated DNA Technologies (IDT). We assembled the gBlocks Gene Fragments together with PGK promoter into sleeping beauty vector, which retains transposase-mediated integration capacity, by following the instructions from Gibson cloning kit (NEB; E2611). All sgRNAs used for hydrodynamic injection were cloned into px330 backbone (Addgene; 42230). Primers and oligos used for cloning are listed in Supplementary Table 1.

### Cell culture

We cultured KPC cells derived from pancreatic cancer model (Kras^G12D^;p53^-/-^) with DMEM (Corning; 10-13-CV) supplemented with 10% FBS (Thermo Fisher Scientific; 16000044). For transfection, we used lipofectamine 2000 reagents (Invitrogen; 11668027) according to the manufacturer’s instructions. At two days post-transfection, we collected cells to isolate genomic DNA (Roche; 06650767001) and total RNA (Qiagen; RNeasy Mini kit, 74104) by following manufacturers’ instructions.

### Hydrodynamic tail vein injection

To prepare large quantities of plasmids, we cultured 250 ml *Escherichia coli* bacteria bearing transformed plasmids and isolated the plasmids using Qiagen Maxi-Prep Endotoxin-free kit (Qiagen; 12362) according to the manufacturer’s instructions. Hydrodynamic tail vein injection was performed as previously described (2). Briefly, we mixed 40 to 60 μg plasmids in 2 ml 0.9% sterile saline solution (w/v) at room temperature and delivered the mixture to mice via tail vein injection within 5-7 seconds. Mice were then warmed by heating pad for 30 min to recover from the injection shock.

### Histology and Immunohistochemistry

We euthanized the mice, collected the tissues, and fixed them overnight in 4% formalin (v/v) overnight. Fixed tissues were then switched to 70% ethanol (v/v) and submitted to the Histology Core Facility of Cold Spring Harbor Laboratory for paraffin embedding and sectioning, and hematoxylin and eosin (H&E) staining. From the unstained slides, we performed immunohistochemistry (IHC). Briefly, we deparaffinized the unstained slides in 100% xylene for three times and hydrated the slides in serial of ethanol (v/v) 100%, 95%, 90% and 70% three times each. We then boiled the slides for 10 min in 1 mM citrate buffer (w/v) with pH 6.0 to retrieve the antigen. We used ImmPRESS Excel Amplified HRP Polymer Staining kit (Vector Laboratories; MP-7601) for the IHC staining. Specifically, BLOXALL reagent was used to block endogenous Horseradish peroxidase (HRP) signal, and 2.5% normal horse serum (w/v) was used to block non-specific binding. Primary antibodies against β-catenin (Santa Cruz; sc-7199), YAP (Cell Signaling Technology; 14074) or Ki67 (Thermo Fisher Scientific; MA5-14520) was diluted 1:200 in PBST buffer and incubated with slides overnight at cold room. We next followed the ImmPRESS kit to amplify the antibody signal by adding the secondary and tertiary antibodies. To visualize the brown color signal, we added 3,3’– Diaminobenzidine (DAB) substrate provided in the kit (Vector Laboratories; MP-7601). To visualize the red color signal, we added biotinylated secondary antibody and used ABC (Avidin/Biotin) system provided in Vectorstain ABC-AP kit (Vector Laboratories; AK5200) and Vector Red substrate kit (Vector Laboratories; SK-5100). The slides were then counterstained with hematoxylin for 2 min, and dehydrated in serial of ethanol 70%, 90%, 95% and 100% three times each. Slides were then immersed in 100% xylenes three times and sealed for long term storage. Images were captured using Olympus microscope (DP72).

### RT-PCR, TOPO cloning and sanger sequencing

We reverse-transcribed isolated mRNA using High-capacity cDNA reverse transcription kit (Thermo Fisher Scientific; 4374966). We then performed LaTaq PCR (Takara, RR002A) to amplify the β-catenin transcripts, and run on 1.5% agarose gel to visualize the amplicon. Skipping transcripts were gel purified using QIAquick gel extraction kit (Qiagen; 28704), cloned to TOPO vector using TOPO TA cloning kit (Invitrogen; 450071) for sanger sequencing (Eurofins Company sequencing service).

### Western blotting

We homogenized the collected tissues in RIPA lysis buffer (CST; 9806) (25 mM Tris-HCl, pH 7.6, 150 mM NaCl, 1% (v/v) NP-40, 1% sodium deoxycholate, and 0.1% (w/v) SDS) containing 1× E-64 protease inhibitor (Sigma; E3132) and centrifuged at 12,000 rpm to obtain clear supernatant. We next quantified the protein concentration of samples using Pierce BCA protein assay kit (Thermo fisher scientific; 23225). Samples were boiled in 4X NuPAGE LDS sample buffer (Invitrogen, NP008) containing 100mM DTT. We run 20 μg total proteins on 4-12% NuPAGE Bis-Tris mini protein gels (Invitrogen; NP0321BOX). Proteins separated by their approximate sizes on the gel were then transferred onto nitro-cellular membrane (Thermo Fisher Scientific; 88018). We next blocked the membrane in blocking buffer (Li-Cor; 927-60001) for 1 h at room temperature and incubated with primary antibodies at 1:2000 dilution against β-catenin Cell Signaling Technology; 8480S), and HSP90 (BD biosciences; 610418) overnight at cold room. Next day, primary antibody signal was detected by secondary antibody IRDye 800CW (Li-Cor; 926-32213) or 680RD (Li-Cor;926-68072)) and visualized using Li-Cor Odyssey imaging machine.

### RNA sequencing

We isolated total RNA from tumors using MirVana miRNA Isolation kit (Invitrogen; AM1561) and removed ribosomal RNA (rRNA) using 186 oligos as described in previously established protocols (50, 51). To enrich RNA >200 nt and remove tRNAs, we purified RNA samples using RNA Clean & Concentrator-100 (Zymo; R1019). We prepared strand-specific libraries for high throughput RNA sequencing (RNA-seq) as described previously (50). All reagents related to RNA-seq library preparation were listed in the Supplementary Table 4. Prepared RNA libraries were sequenced by 79 + 79 paired-end reading using NextSeq550 (Illumina).

Raw reads were processed using DolphinNext (52) RNA-Seq pipeline Revision 2. STAR (version 2.6.1)(53) and RSEM (version 1.3.1) (54) indices were created based on the mouse genome assembly mm10 and gene annotations downloaded from UCSC genome browser. RSEM software package (54) was then used to estimate relative gene expressions using STAR (53) aligner. To examine differentially expressed transcripts, we used DEBrowser (55), interactive differential expression analysis tool. We compared the raw gene counts estimated by RSEM from biological replicates in different conditions using DESeq2 (56). We filtered out genes whose max log10 raw counts less than 1. Gene ontology (GO) term enrichment analysis was performed using DAVID Functional Annotation Tool (57) The complete list of GO term categories with significant enrichment was extracted. Gene Sets Enrichment Analysis was conducted using GSEAPreranked (58) software package on differential analysis results generated by DESeq2.

### Exon skipping events in human tissue and cancer samples

#### The Cancer Genome Atlas (TCGA) samples

Aligned Genomic Bam RNAseq files (GRCh38/hg38 coordinates) for TCGA tissue samples with primary diagnosis ‘Hepatocellular carcinoma, NOS’ (HCC) where downloaded from Genomic Data commons TCGA-LIHC (dbGAP accession phs000178.v11.p8, project ID 26811). As described in the TCGA data processing protocol (https://docs.gdc.cancer.gov/Data/Bioinformatics_Pipelines/Expression_mRNA_Pipeline/), genomic Bam RNAseq files were generated using STAR (53) in two-pass mode with splice junction detection step, allowing for downstream splicing analysis. We used *samtools* (59) to extract all reads mapping to the coding region of *CTNNB1*: ‘samtools view -o output.bam input.bam chr3:41194741-41260096’.

#### ExonSkipDB

We queried ExonSkipDB (41) for beta-catenin (*CTNNB1*) exon skipping events in any tissue or cancer as curated based on data from the genotype-tissue expression (GTEx) portal and TCGA. ExonSkipDB reports its exon skipping events in GRCh37/hg19 coordinates. We used the UCSC *liftover* tool (60) with the following parameters to convert the coordinates of the exon skipping events to GRCh38/hg38: Minimum ratio of bases that must remap: 0.95; Allow multiple output regions: off; Minimum hit size in query: 0; Minimum chain size in target: 0; If thickStart/thickEnd is not mapped, use the closest mapped base: off. All originally reported positions were successfully converted.

#### Visualization of TCGA and ExonSkipDB events

Exon skipping events identified via ExonSkipDB and TCGA HCC RNAseq reads were visualized using *Gviz* (61). As reference, the exon and intron position for canonical beta-catenin isoforms expressed in liver (ENST00000349496, ENST00000396185, ENST00000396183) were added as a separate track.

## Acknowledgements

We thank CSHL Cancer Center Shared Resources (Animal, Flow Cytometry and Histology, Sequencing and Bioinformatics Core Facilities) supported by NCI Cancer Center Support grant 5P30CA045508. We thank Soren Hough (Cambridge University, UK) and Angela Park (Weill Cornell Medical College, USA) for editing of the manuscript. This work was supported by the National Institutes of Health (P30CA045508-33 to S.B.), Mark Foundation for Cancer Research (20-028-EDV to S.B. and T.J.) The Oliver S. and Jennie R. Donaldson Charitable Trust (S.B.), STARR Cancer Consortium (I13-0052, S.B.). A.K. was supported by the National Center for Advancing Translational Sciences grant #UL1 TR001453–01. D.M.O was supported by the Swedish Research Council grant 2020-03818.

## Authors’ contributions

H.M., D.M.O, and S.B. conceived the study. H.M. and J.Y., Z.W., Y.G. performed experiments. A.K. H.V.M and Y.J. performed bioinformatic analysis. J.E.W. performed histopathological analysis. H.M., D.M.O, and S.B. wrote the manuscript with comments from all authors.

## Data Availability

Sequencing data are available from the National Center for Biotechnology Information Sequence Read Archive using accession number GSE164109.

---

**Supplementary Figure S1.**
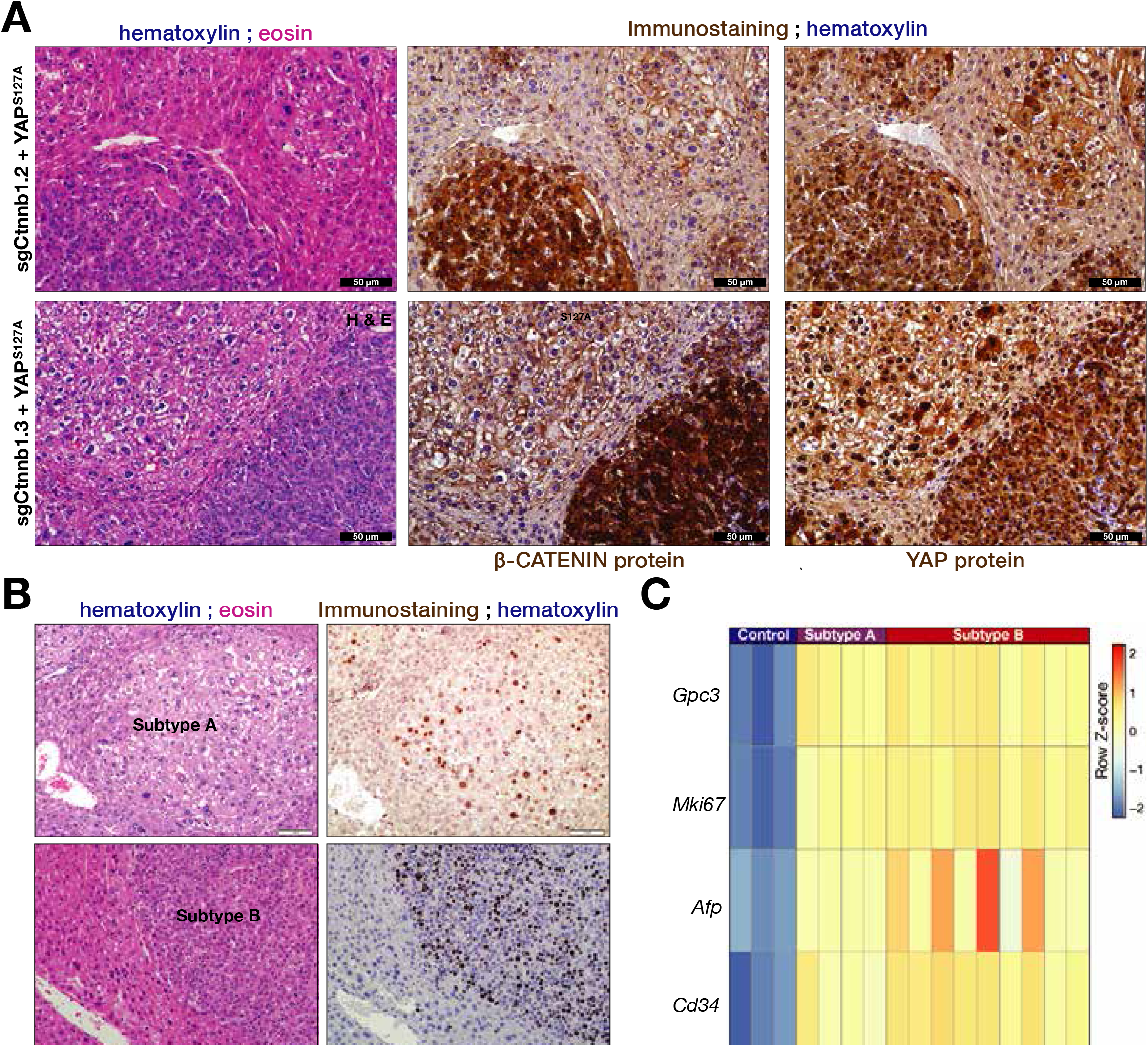
Tumors driven by sgCtnnb1.2/YAP^S127A^, sgCtnnb1.3/YAP^S127A^. (**A**) Hematoxylin and eosin (H&E) staining shows the histology and IHC shows the β-catenin and YAP staining from tumors driven by sgCtnnb1.2/YAP^**S127A**^ or sgCtnnb1.3/YAP^**S127A**^. Brown color indicates positive IHC signal. (**B**) H&E and Ki67 IHC staining from the serial sections of Subtype A and B tumors. Brown color indicates positive Ki67 IHC signal. (**D**) Heat-map shows liver tumor marker genes, *Gpc3, Mki67, Afp*, and *Cd34* in Subtype A and B.

**Supplementary Figure S2.**
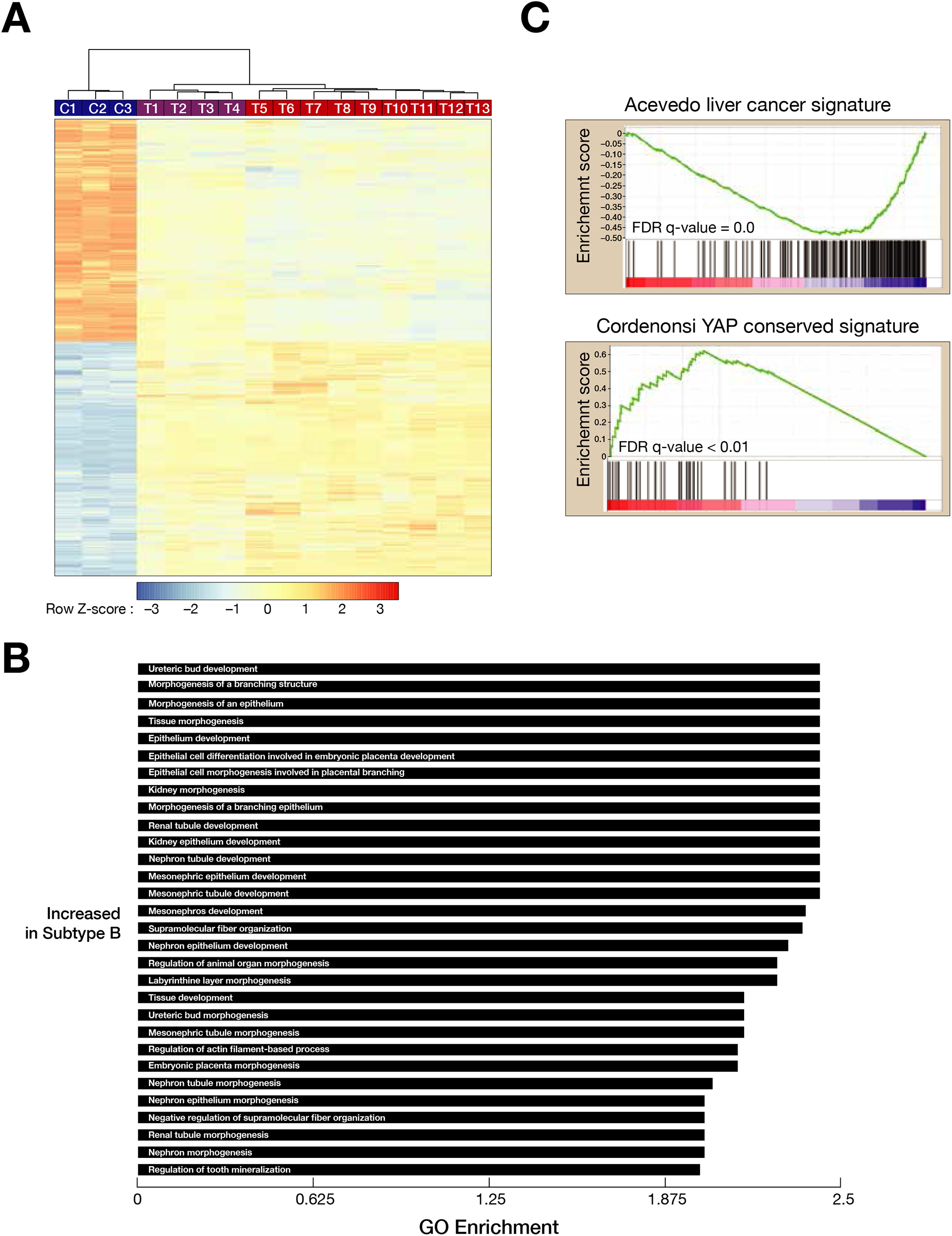
Characterization of tumors driven by sgCtnnb1.1/YAP^S127A^. **(A)** Unsupervised hierarchical clustering shows distinct transcriptome signatures of tumor samples and their corresponding controls. C denotes control, T stands for tumor. **(B)** Gene Sets Enrichment Analysis (GSEA) shows the enrichment of tissue development and morphogenesis genes in Subtype B. (**C**) Both Subtype A and B show liver cancer and Yap activation signature.

**Supplementary Figure S3.**
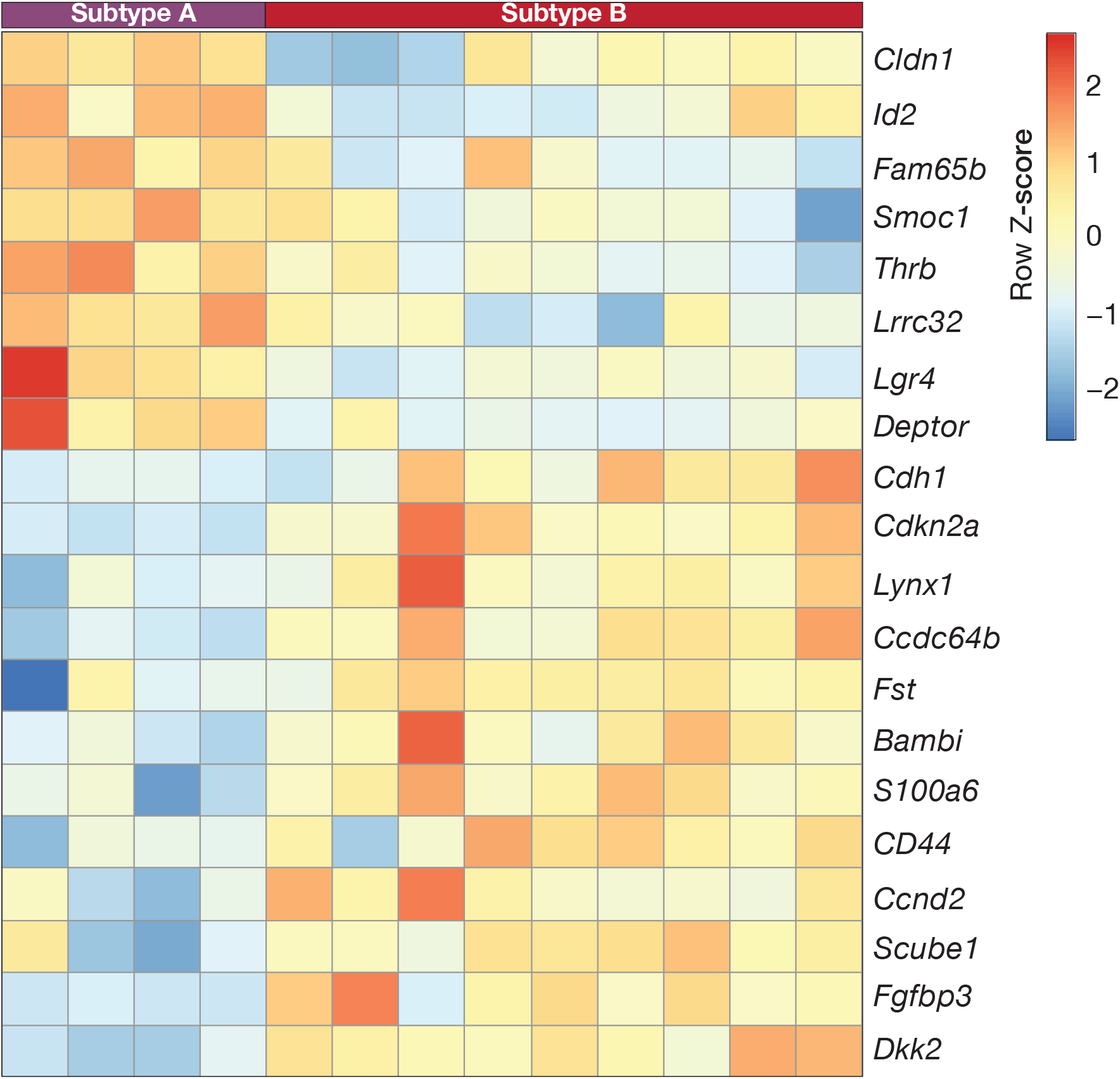
Expression pattern of β-catenin target genes in Subtype A and B tumors. Heatmap shows the expression of selected representative β-catenin target genes in Subtype A and B.

**Supplementary Figure S4.**
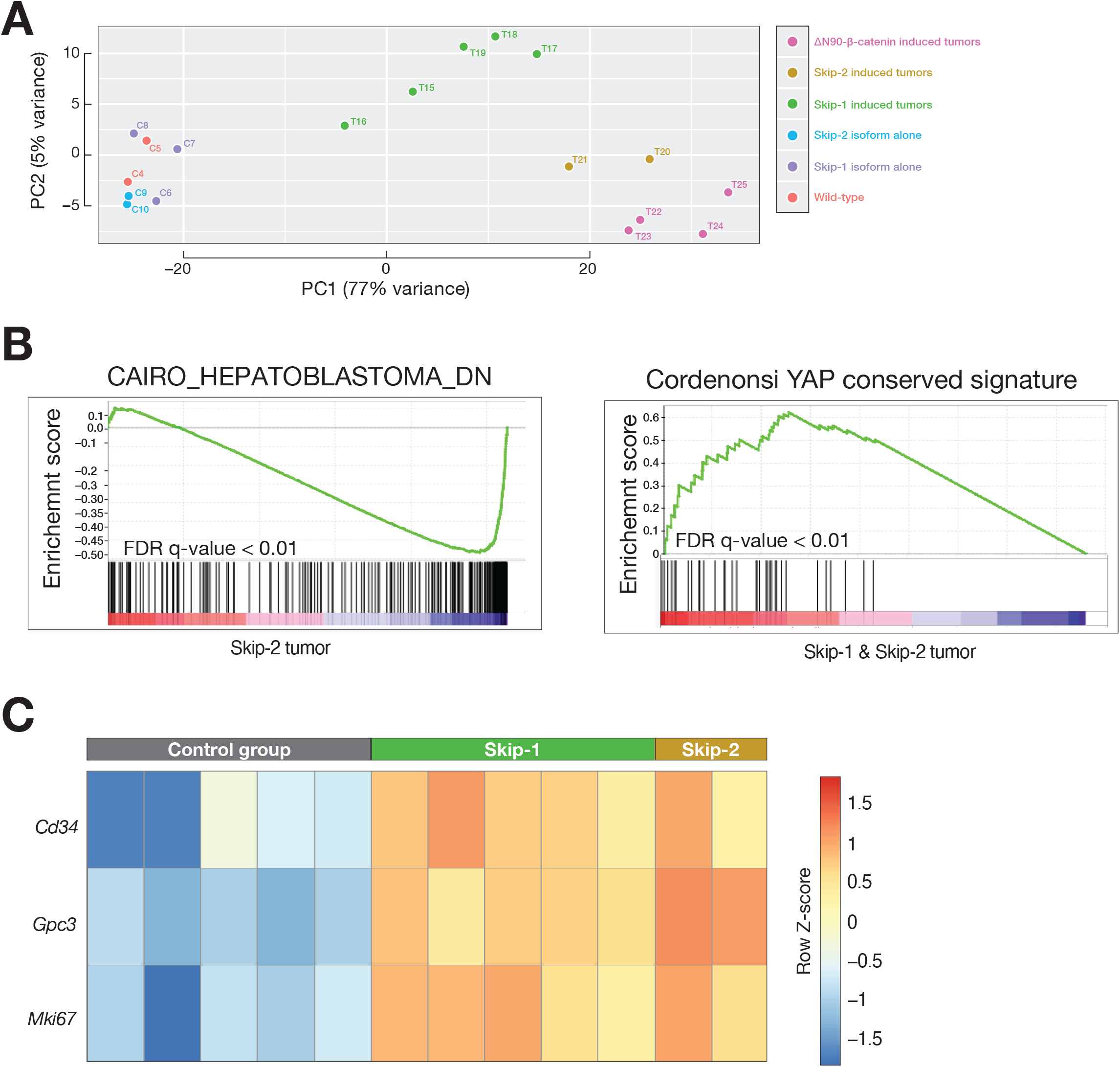
Skip-2 tumors show hepatoblastoma feature. (**A**) Principal Component Analysis (PCA) groups samples based on their transcriptome signature. (**B**) Gene Sets Enrichment Analysis (GSEA) shows Hepatoblastoma signature in Skip-2 tumors. Both Skip-1 and Skip-2 show YAP activation signature. (**C**) Heatmap shows the expression of liver tumor marker genes, *Cd34, Gpc3* and *Mki67*, in Skip-1 and -2 tumors compared to corresponding controls.

**Supplementary Figure S5.**
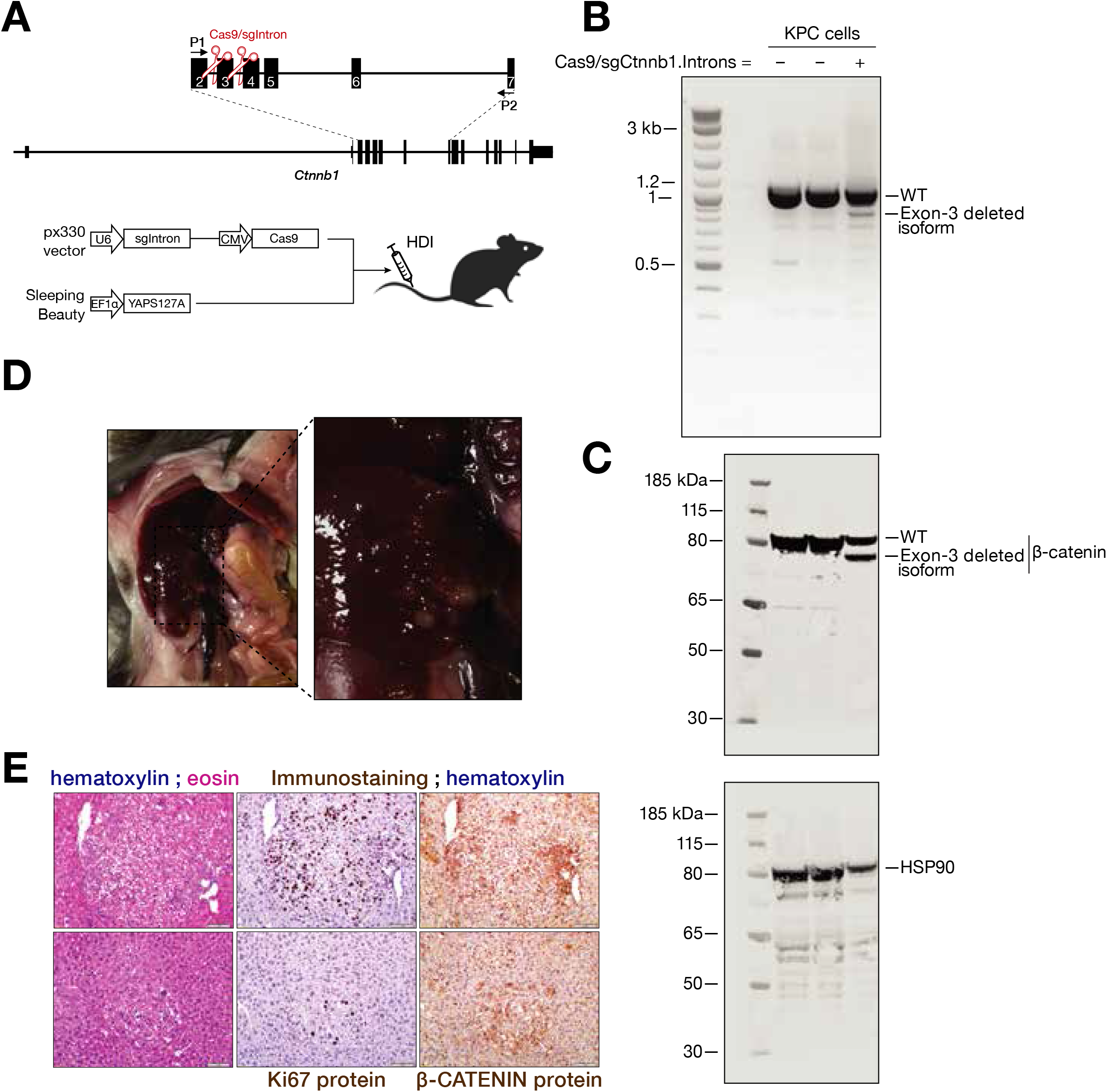
Genomic deletion of Exon 3 of *Ctnnb1* (β-catenin) leads to tumor formation in synergy with YAP^S127A^. (**A**) Schematic representation of HDI delivering two sgRNAs that target introns flanking Exon 3 of β-catenin and SB-YAP^S127A^. P1 and P2 Primers are used to detect deletion of Exon 3. (**B**) RT-PCR analysis detects the transcripts with Exon 3 deletion using P1/P2 primers from the total RNA of Pancreatic Kras^G12D^; p53^-/-^; Cre+ (KPC) cells transfected with two sgRNAs targeting introns as illustrated in (A). (**C**) Western blotting detects β-catenin protein isoform without Exon3 in the protein lysate from KPC cells transfected with sgRNAs targeting introns as shown in (A). (**D**) Image captured by iPhone camera shows tumor formation driven by sgRNAs that target introns flanking Exon 3 of Ctnnb1 in synergy with YAP^S127A^. (**E**) Hematoxylin and Eosin (H&E) staining shows the histology of tumors while IHC shows Ki67 and β-catenin staining in tumor induced by two sgRNAs targeting introns flanking Exon3 in combination with YAP^S127A^.

**Supplementary Figure 6.**
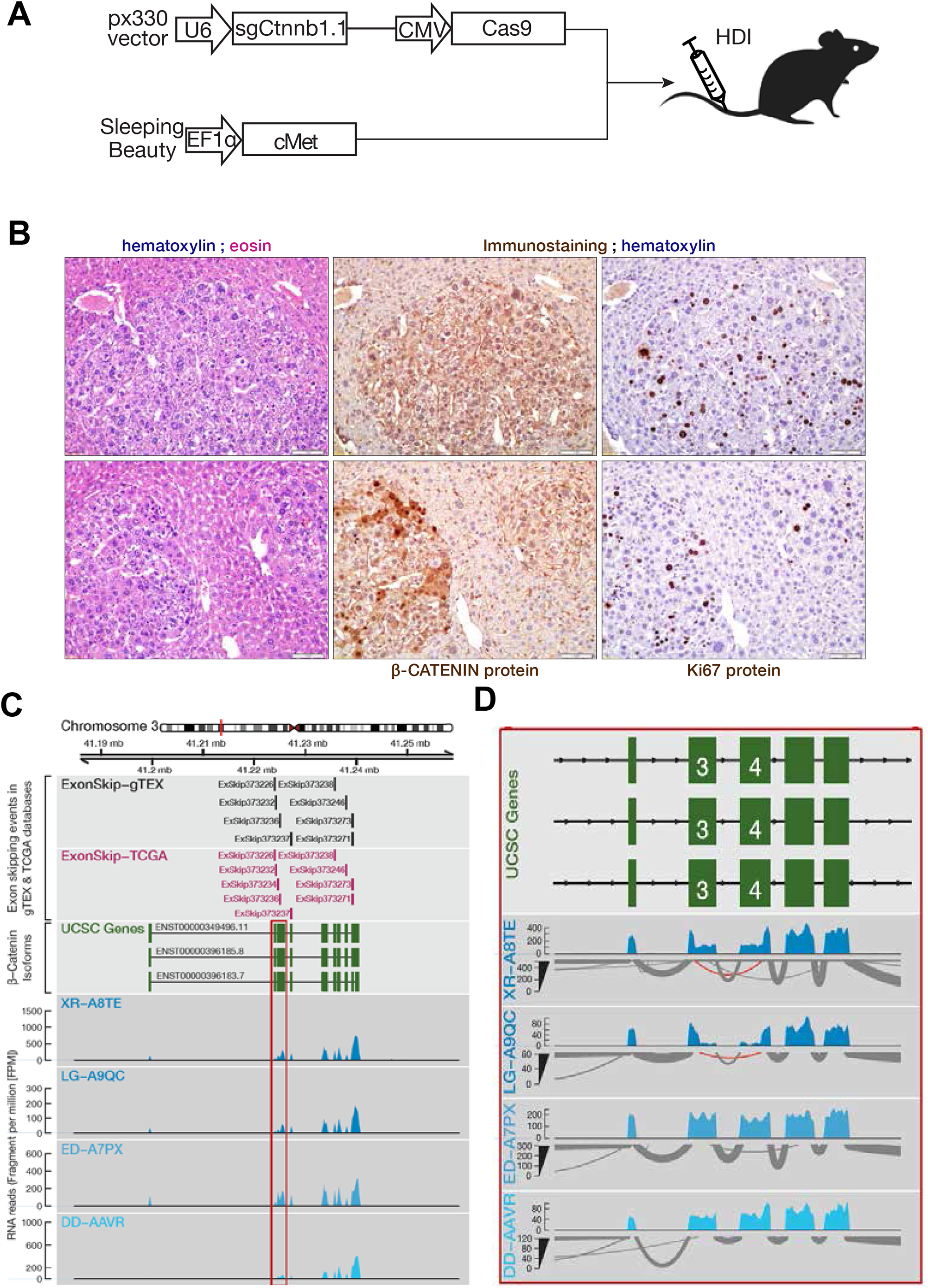
Exon 3 and 4 skipped β-catenin transcript presents in human hepatocellular carcinoma patients. (**A**) Schematic representation of HDI delivering the oncogene c-Met along with sgCtnnb1.1. (**B**) H&E staining and IHC staining of Ki67 and β-catenin from the serial sections of sgCtnnb1.1/c-Met induced tumors. Brown color indicates positive IHC signal. (**C**) UCSC genome browser view of *CTNNB1* gene with RNA sequencing reads from hepatocellular carcinoma patients obtained from The Cancer Genome Atlas (TCGA). (**D**) Exon-skipping events over the exons of *CTNNB1* gene identified via ExonSkipDB (41).

